# Prots2Net: a PPIN predictor of a proteome or a metaproteome sample

**DOI:** 10.1101/2022.06.24.497208

**Authors:** Adrià Alcalá, Mercè Llabrés

## Abstract

**Motivation:** All molecular functions and biological processes are carried out by groups of proteins that interact to each other. Proteins interactions are modeled by simple networks called Protein-Protein Interaction Networks (PPINs) whose nodes are proteins and whose edges are the protein-protein interactions. PPINs are broadly accepted to model the protein’s functional relations, and their analysis has become a key ingredient in the study of protein functions. New proteins are collected every day from metaproteomic data, and their functional relations must be obtained with high-throughput technology. Retrieving protein-protein interaction data experimentally is a very high time-consuming and labor-intensive task. Consequently, in the last years, the biological community is looking for computational methods to correctly predict PPIs.

**Results:** We present here Prots2Net, a tool designed to predict the PPIs of a proteome or a metaproteome sample. Our prediction model is a multilayer perceptron neural network that uses protein sequence information only from the input proteins and interaction information from the STRING database. To train the model, Prots2Net explores the PPIs retrieved from the STRING database of two selected species. The tests, reported here on the Yeast and the Human datasets, show that Prots2Net performs better than the previous prediction methods that used protein sequence information only. Therefore, considering the information of PPI data available on the STRING database improves the PPI prediction.

**Availability:** https://github.com/adriaalcala/prots2net

## 1 Introduction

Proteins are the main characters in cellular processes, and their interactions perform, among others, signal transduction, immune response and cellular organization functions. These functions are performed by sets of proteins that interact to each other. In order to understand the mechanics of these biological processes and further discovery of new functions underlying in a biological system, proteins interactions must be detected and analyzed.

Different methodologies to experimentally retrieve protein-protein interaction (PPI) data, such as yeast two-hybrid screens[12, 14], protein chips[34], tandem affinity purificaton[22], immunoprecipitation [19] and spectrometric protein complex identification [9] among others, have been developed in the last decade. As a result, a considerable amount of PPI data has been generated and stored in different databases [26, 29] which allows the study and analysis of PPIs. Such analysis can be performed under different computational techniques, and inference statistics has been proved to be successful at PPI prediction. This fact, together with the acceptance that experimental methods to detect PPIs are too expensive in terms of time and labor, has generated an explosion of computational methods to predict PPIs. These computational methods rely on different machine learning technologies, such as deep neural networks [4], convolutional neural networks [27], rotation forest [28, 16, 20], support vector machine [31, 8, 33, 20], extreme learning machine with principal component analysis [32], *k*-nearest neighbors [30], and random forest [3, 20] among others, and different data types to train the models, such us literature mining knowledge [2], gene fusion [5], phylogenetic profiles [25], gene ontology annotations [17], gene neighborhood [7], and co-evolution analysis of interacting proteins [13]. However, the amount of new sequence data that is created every day and its lack of a priori information, makes very popular those methods that predict PPIs based on protein sequence information [10, 15, 16, 1]. Despite the simplicity of predicting interactions based only on protein sequence information, the results reported by those methods prove that, they perform well and achieve good measures of error prediction [16].

Under the requirement of predicting interactions based only on protein sequence information together with the idea to also exploit the PPIs information stored in the databases, we have designed a PPIs predictor, Prots2Net, based on the protein sequences and two protein-protein interaction networks (PPINs). The training data of our model are two PPINs from the STRING database [26] selected by the user. To build the prediction model, PPIs in one of the selected species are predicted through PPIs of orthologs proteins in the other species, and the PPIs information from the second species is used as the true output in the training. The tests carried on to evaluate the performance of Prots2Net compared to other 12 existing methods, reported values above 95% in accuracy, sensitivity, and precision error measures on the Yeast and Human datasets. Thus improving the results obtained by the others methods.

## 2 Methods

In this section, we present our tool *Prots2Net* to predict PPIs. As shown in Figure 1,Prots2Net is structured in the following steps:

1. Input data (GUI component).
2. Network prediction (Server component).
3. Visualize data (GUI component).

**Figure 1.**
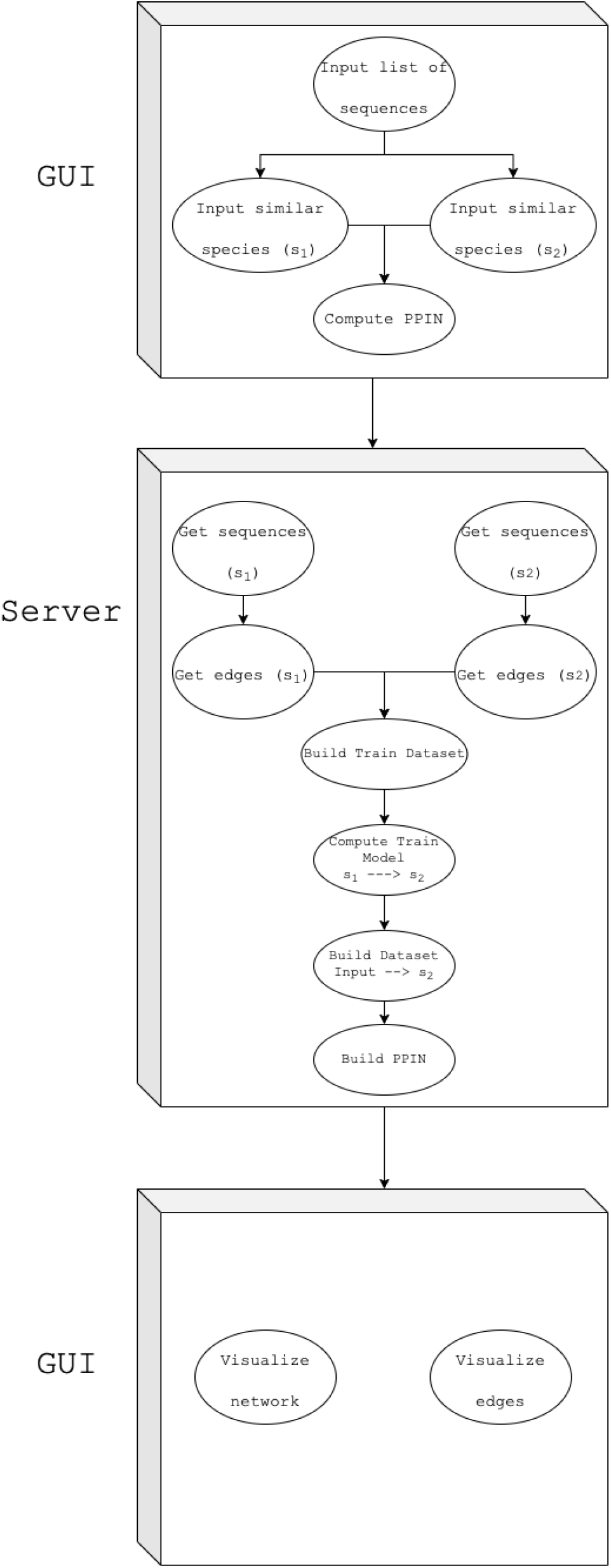
Diagram explaining the workflow of Prots2Net.

In the first stage of Prots2Net, the user inserts the proteins sequences whose PPIs are requested and selects two species from the STRING database used to train the model. In the second stage, Prots2Nets builds the model and predicts the interactions. Finally, the user can visualize and analyze the predicted PPIN in the third stage. Besides this workflow, the code is organized in 2 main components: A backend server, where all the computations are performed and a Graphical User Interface (GUI), where the user interacts.

### Server

The backend server carries on all the computations needed to obtain the PPIs predictions. The workflow of the entire job is as follows: first compute a training model from the two selected species. Then, read the proteins sequences in the input file. And finally, predict the output PPIN using the computed training model.

The overall idea behind this model is that two proteins interact if their orthologs, in another species whose PPIN is known, interact. Thus, to train the model, the user must select two species from the STRING database. Next, the PPIs information from the first species is transferred to the second species, through orthologs proteins, to predict the interactions. Finally, the PPIs information from the second species is used as the true output during training. We explain below the details of the whole process.

### Training model dataset

The training model is constructed using the library sklearn [21] as follows: let *s*_1_ and *s*_2_ be the two species selected by the user from the STRING database, and let *P*_1_ and *P*_2_ be the corresponding set of proteins of *s*_1_ and *s*_2_, respectively. For every pair of proteins, *p*_21_ and *p*_22_ in *P*_2_, Prots2Net takes into account the following information as input to build the training dataset:

- The bitscore *b*_1_ of the sequence alignment between the protein *p*_21_ and its most similar protein *p*_11_ in *P*_1_.
- The bitscore *b*_2_ of the sequence alignment between the protein *p*_22_ and its most similar protein *p*_12_ in *P*_1_.
- The combined interaction score in *s*_1_ between *p*_11_ and *p*_12_ when they interact. If there is no interaction between *p*_11_ and *p*_12_, it is set to 0.

The true output during training, is 1 i.e., *p*_21_ and *p*_22_ are considered to interact, if the combined interaction score in *s*_2_ between *p*_21_ and *p*_22_ is greater than 999. Otherwise, the output is 0 and it is considered that *p*_21_ and *p*_22_ do not interact.

### Neural Network Model

The neural network model used in this tool is an MLP (multilayer perceptron neural network) with 2 hidden layers (see [24] for a brief description of neural networks). As we can see in Figure 2, the input layer has 3 input values which are the values of *b*_1_ and *b*_2_ defined in the training dataset, but considering in this step the pair of proteins *p*_21_ and *p*_22_ from the user’s input set, and the combined interaction score in *s*_1_ when *p*_11_ and *p*_12_ interact, or 0 otherwise. The output layer has only one value, since we are only considering one class, that indicates if *p*_21_ and *p*_22_ interact. As a set of default parameters we considered those listed below, though all of them can be easily changed in the code:

- the activation function used is *ReLU*,
- the optimization algorithm used is *ADAM*.
- the L2 penalty is set to 10^−6^.
- the tolerance is set to 10^−8^, that is, the minimal improvement between steps to continue the training.
- the maximum number of iterations is 30, 000.

**Figure 2.**
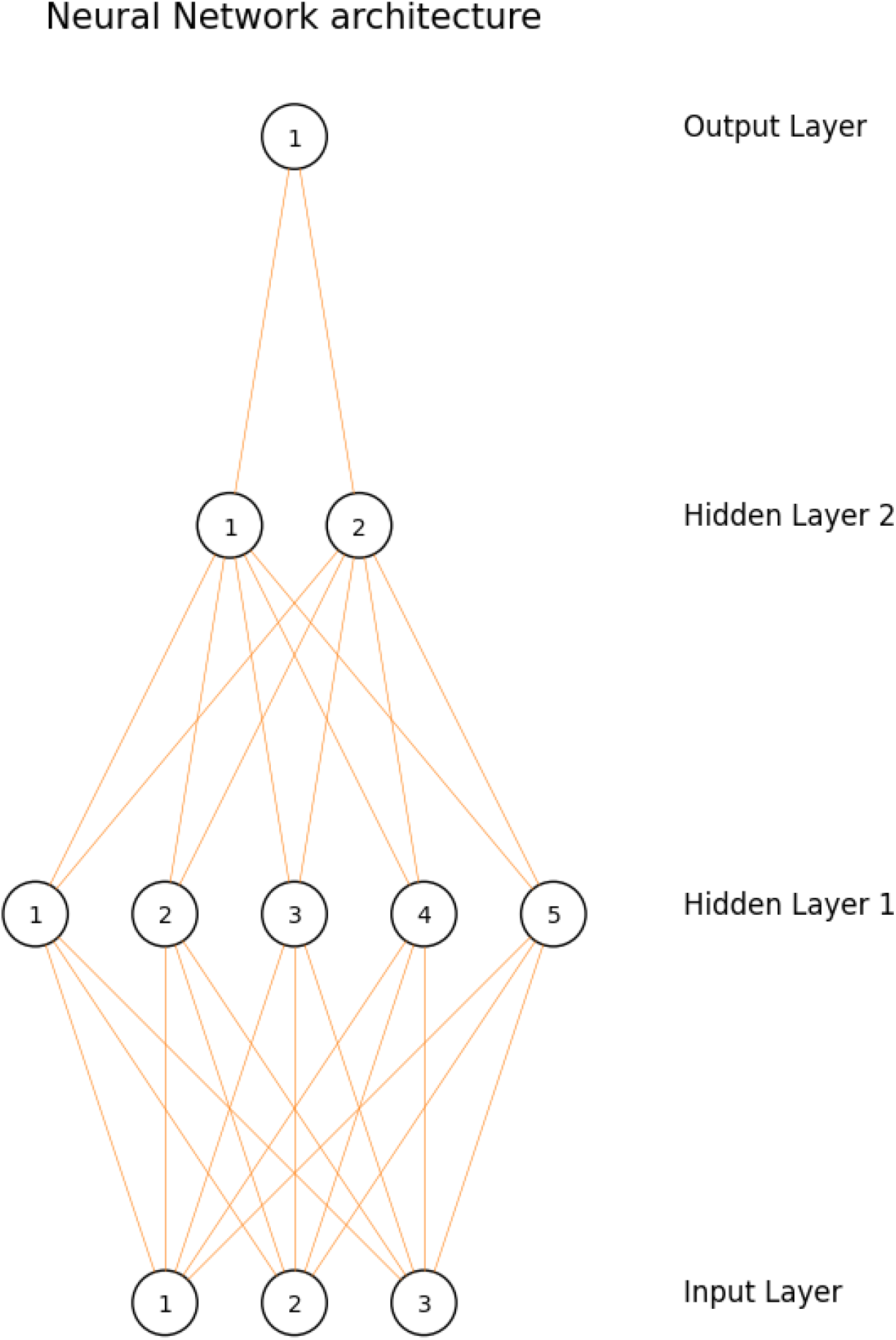
Neural Network architecture.

### Reused data

Generated data for every process is stored in the server under the *data* directory considering the species identifiers. Thus, the following relative paths are created:

- *sequences*, a folder that contains all the sequences files.
- *blast*, a folder that contains the results of the blast computations. As an example, if the user’s main species identifier is *main-species* and *species1* and *species2* are the selected two similar species. Then, the stored files are *blast/result_species1_species2*.*csv* and *blast/result_main-species_species2*.*csv*.
- *edges*, a folder that contains the edge list of every network used in the training step. Notice that these files are the list of edges downloaded from the STRING database under the edge score threshold.
- *nets*, a folder that contains all predicted networks. For instance, in the previous example, the stored file is *nets/Net_main-species_species2*.*csv*.
- *graphs*, a folder that contains the images to visualize all predicted networks. Following the previous example, the stored is *data/graphs/Net_main-species_species2*.*png*.
- *models*, a folder that contains the trained models. Following the previous example, the stored file is *models/Model_species1_species2*.*pickle*.

Since all stored files have their names referred to the species identifiers, all generated models are reused when the selected species have been already considered. Hence, we recommend the user to take this into account when selecting the identifiers names.

### Graphical User Interface (GUI)

This GUI (Figure 3) is implemented in Python [6] and use tkinter libraries (https://docs.python.org/3/library/tkinter.html) to allow the users to interact with the application. This interface has 4 different tabs:

1. A tab to insert the proteins sequences.
2. A tab to visualize the predicted network.
3. A tab to obtain the list of nodes (proteins).
4. A tab to obtain the list of edges (interactions).

**Figure 3.**
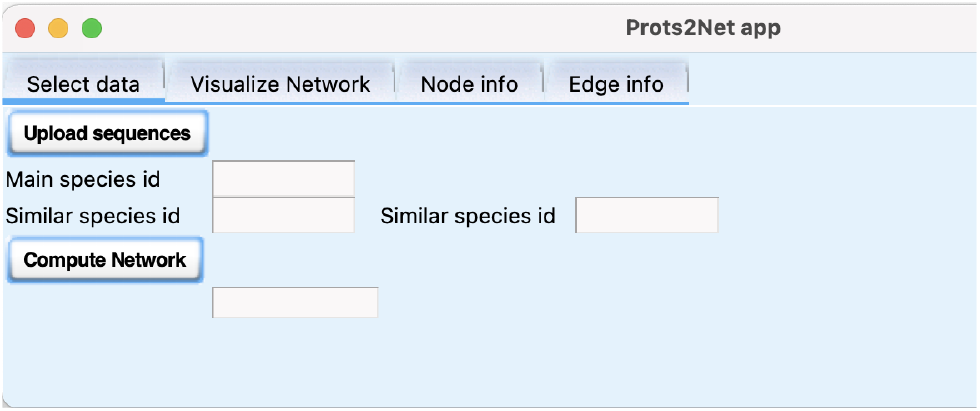
Graphical user interface of Prots2Net.

### Insert data tab

In this tab, the protein sequences are inserted to predict their PPIs. This tab is composed by the following components:

- A button to select a file with the input data.
- 3 text boxes to introduce the species identifiers.
- A button to compute the network.

The user must click the *Upload sequences* button to select a file with the protein sequences in FASTA format, using a file selector. Once the user has uploaded the protein sequences, an identifier for this file, which we call the main species identifier, must be introduced. Next, the user must select 2 (similar) species from the STRING database and finally click the Compute Network button to predict the PPIs.

### Visualize network tab

The user can visualize the predicted PPIN by clicking on the *Plot graph* button and selecting the predicted network file. The image file is stored in the relative path *data/graphs*. This visualization is of use when the network is not so huge, as one we can see in Figure 4 and Figure 5.

**Figure 4.**
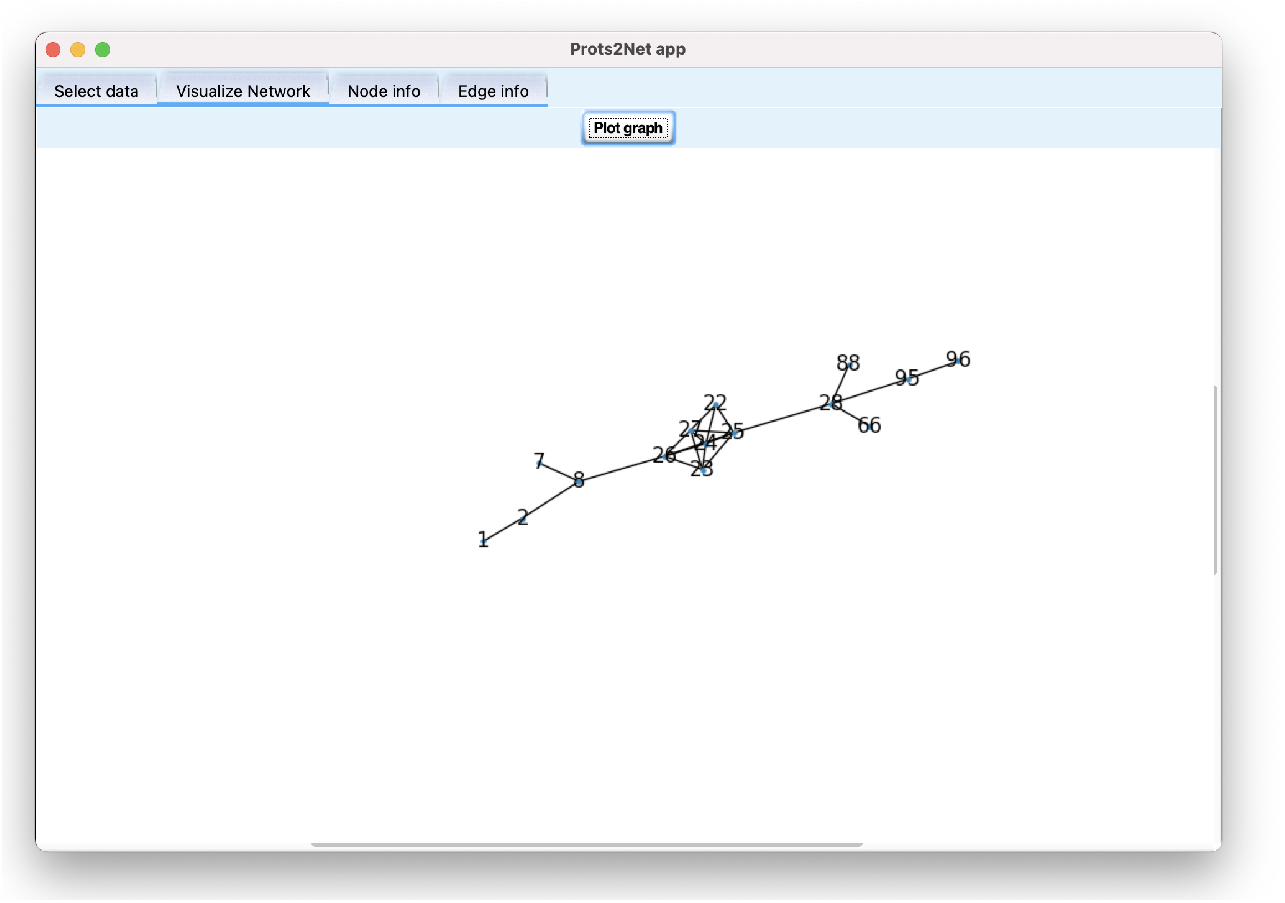
Small network visualization in Prots2Net app.

**Figure 5.**
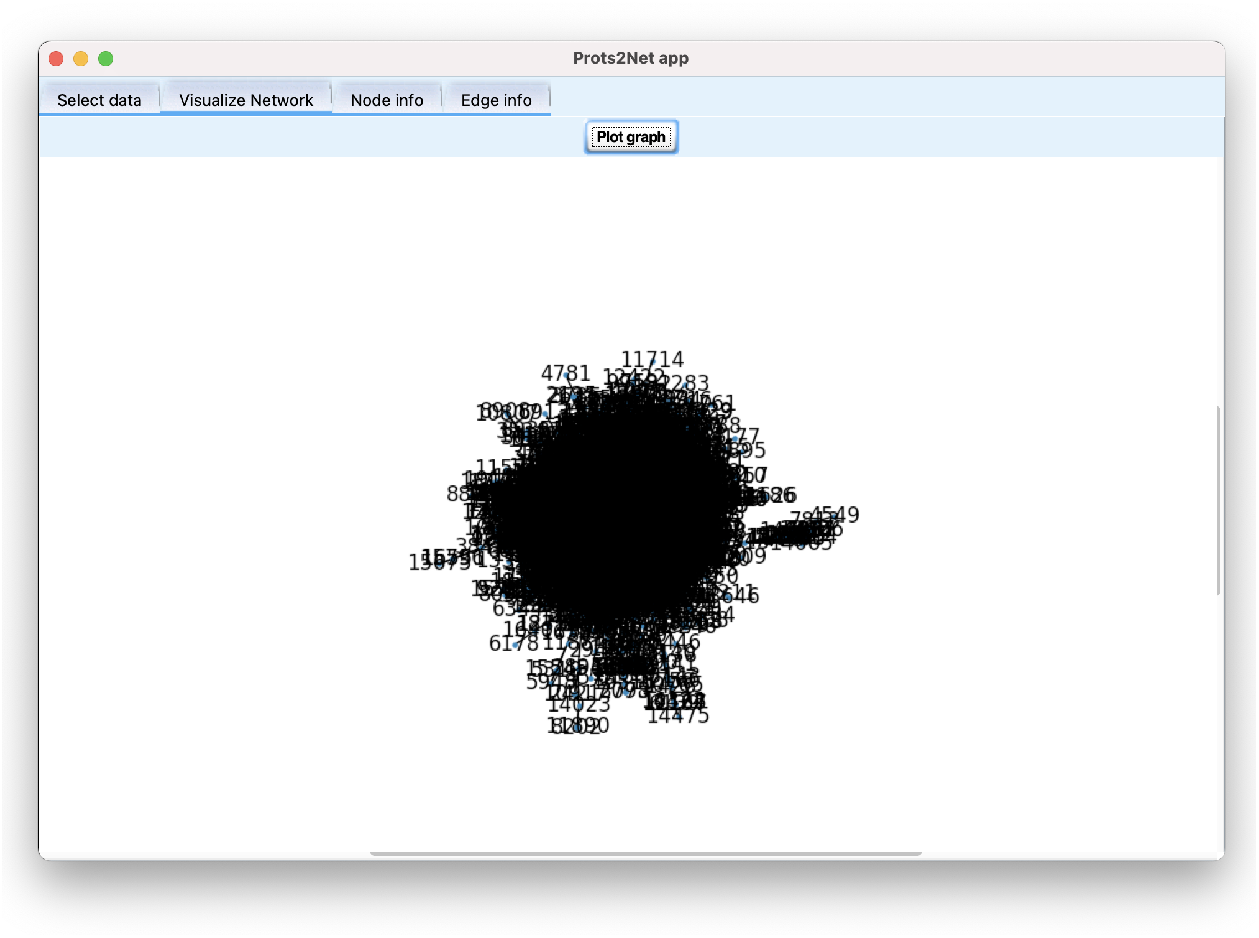
Huge network visualization in Prots2Net app.

The list of nodes and edges of the predicted network are provided by clicking on the *Node info* and *Edge info* buttons, located on top of the window.

## 3 Results and Discussion

We developed the tool Prots2Net that implements the methodology proposed here. The tool was implemented in Python and, it is available at https://github.com/adriaalcala/prots2net. In order to evaluate Prots2Net, and compare our tool with other existing tools, we performed three tests to predict the PPIs in species from different kingdoms. The species that we considered are the following:

1. to predict the PPIs in the Rhodothermus profundi (NCBI 633813), we selected the bacteroidetes Rhodothermus marinus (NCBI 518766) and Salinibacter ruber (NCBI 309807) as similar species.
2. to predict the PPIs in Saccharomyces cerevisiae (NCBI 4932), we selected Millerozyma farinosa (NCBI 4920) and Naumovozyma dairenensis (NCBI 27289) as similar species.
3. to predict the PPIs in Homo sapiens (NCBI 9606), we selected Pan paniscus (NCBI 9597) and Gorilla gorilla (NCBI 9593) as similar species.

Notice that, in every test we selected, as similar species, two species that are taxonomically close to the species whose PPIs are predicted. More precisely, in the first test, we consider as similar species another two bacteroidetes. In the second test, we selected as similar species another two fungi and, in the third test, we considered another two hominidae.

As explained in the previous section, we considered as parameters, a multilayer perceptron classifier (MLP) model with two hidden layers with 5 and 2 nodes respectively and the configuration explained in the Methods section (see Neural Network Model subsection). Then, for every test, we applied Prots2Net to the input protein sequences considering the mentioned similar species. To validate the proposed model, we considered the following error measures:

- Accuracy: defined as 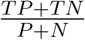,
- Precision: defined as 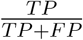,
- MCC: the Matthews correlation coefficient measure is defined as 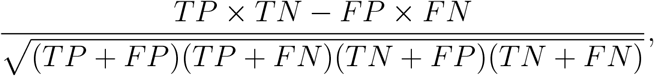,
- Sensitivity: defined as 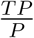,
- Specificity: defined as 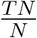,
- FallOut: defined as 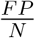,
- F1-score: defined as 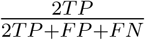

where *TN* is the number of true negatives, i.e., the non-interacting proteins that are predicted correctly; *TP* is the amount of true positives, i.e., the interacting proteins that are predicted correctly; *FN* is the number of false negatives, i.e., the interacting proteins that are predicted to be non-interacting; and *FP* is the amount of false positives, i.e., the non-interacting proteins that are predicted to interact. Additionally, the receiver operating characteristic (ROC) curves and the area under the ROC curve (AUC) were also calculated to further evaluate the discriminatory accuracy of the proposed model. In order to obtain a highly reliable dataset as true PPINs, we considered for the selected species the corresponding PPINs from the STRING database with a combined interaction score threshold of 999, and a set with the same size of non-interacting proteins. That is, there is a true interaction between two proteins when their combined interaction score is equal or greater to 999.

To contextualize the results obtained with Prots2Net, and compare them with the currently known methods, we considered the method introduced in [16] as well as all the methods reported there. Hence, we considered our results with the results obtained by 12 other computational methods.

### 3.1 Test 1. Rhodothermus profundi predictions

Figure 6 shows the evaluation results obtained by Prots2Net in the first test. As we can observe, the *precision* and *accuracy* values are very high, almost 1, and the *Fall-out* values are nearly 0. This is a consequence of the fact that the number of false positives is almost 0, as we can observe in Figure 7. Indeed, we can see there that there is only 1 false positive, when the threshold is 0.5, i.e., there is only one interaction that Prots2Net predicts, and it is wrong. Notice that, even with this high precision, there is no loss of accuracy, which is higher than 0.91. This excellent performance is also displayed in the ROC curve, in Figure 8, where we can observe that the ROC curve has a high *AUC* value of 0.9261.

**Figure 6.**
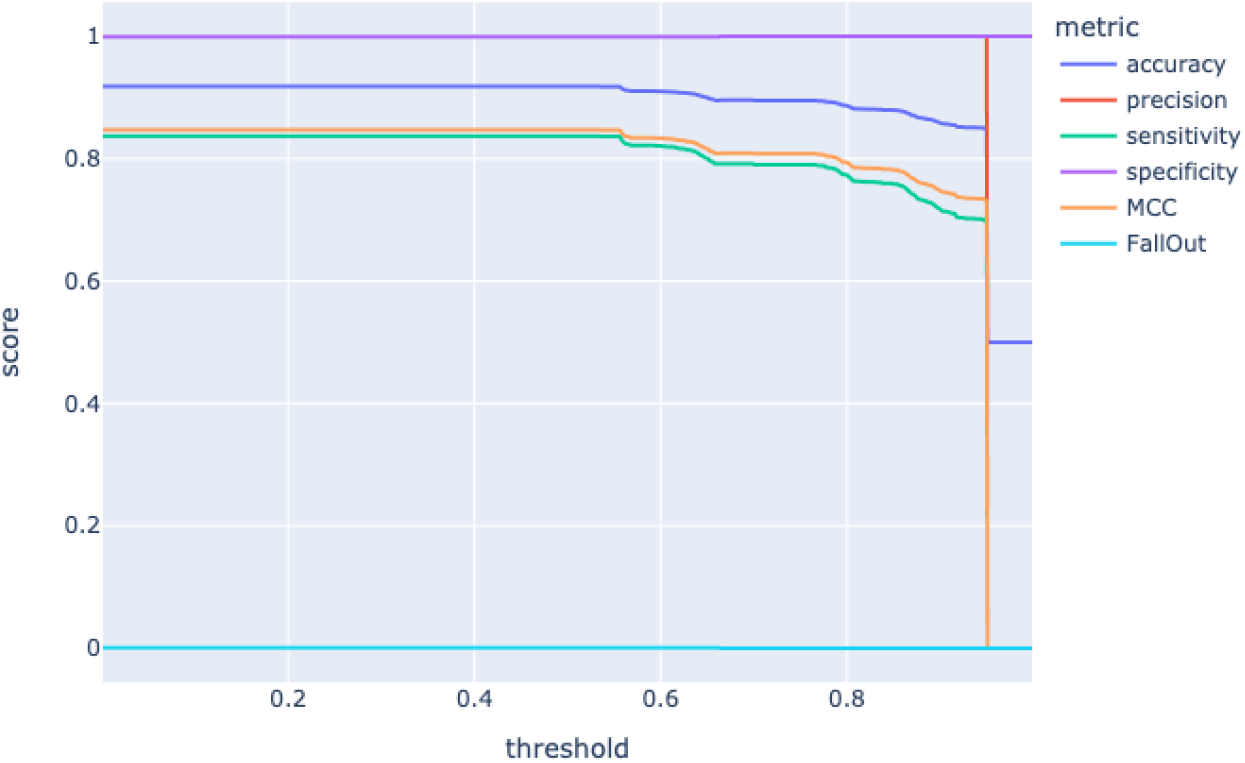
Metrics for Rhodothermus profundi PPIs prediction.

**Figure 7.**
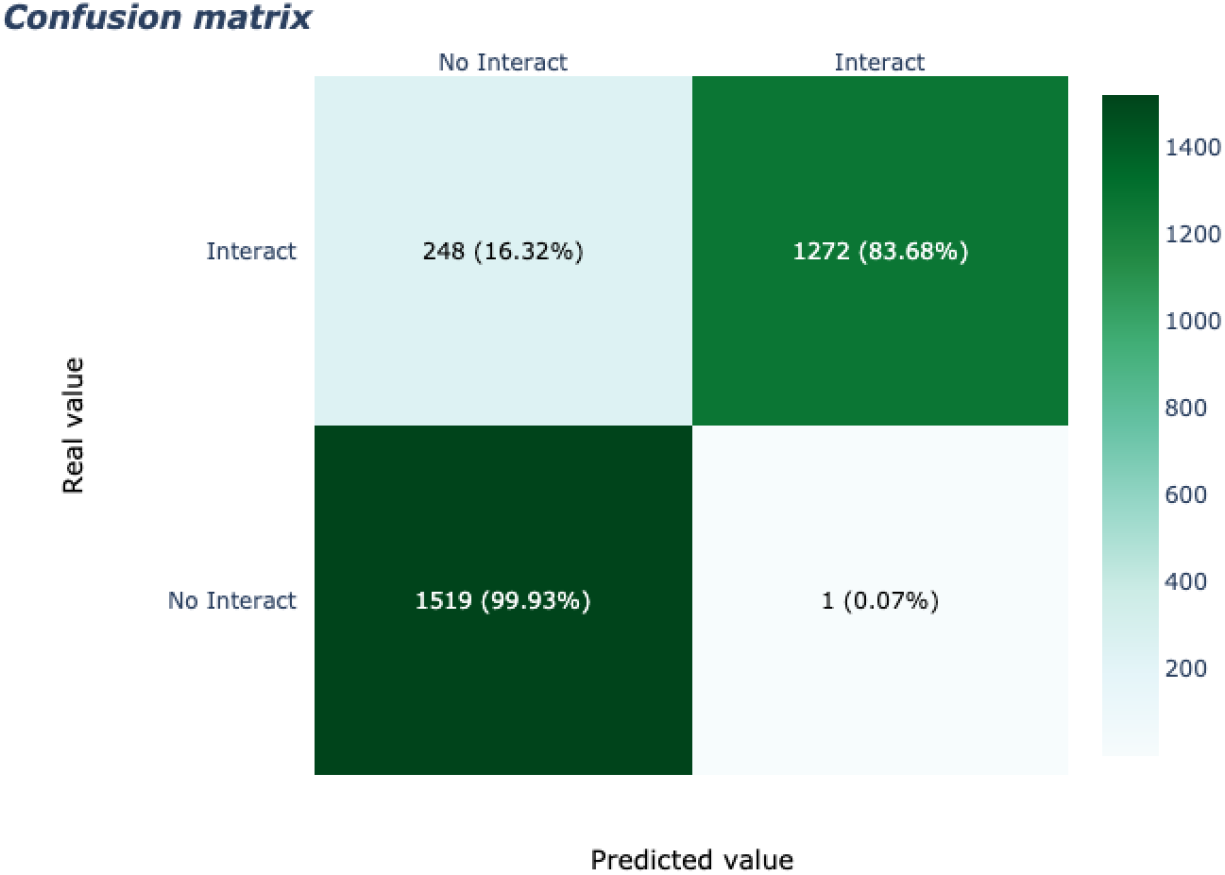
Confusion matrix for Rhodothermus profundi with a threshold of 0.5.

**Figure 8.**
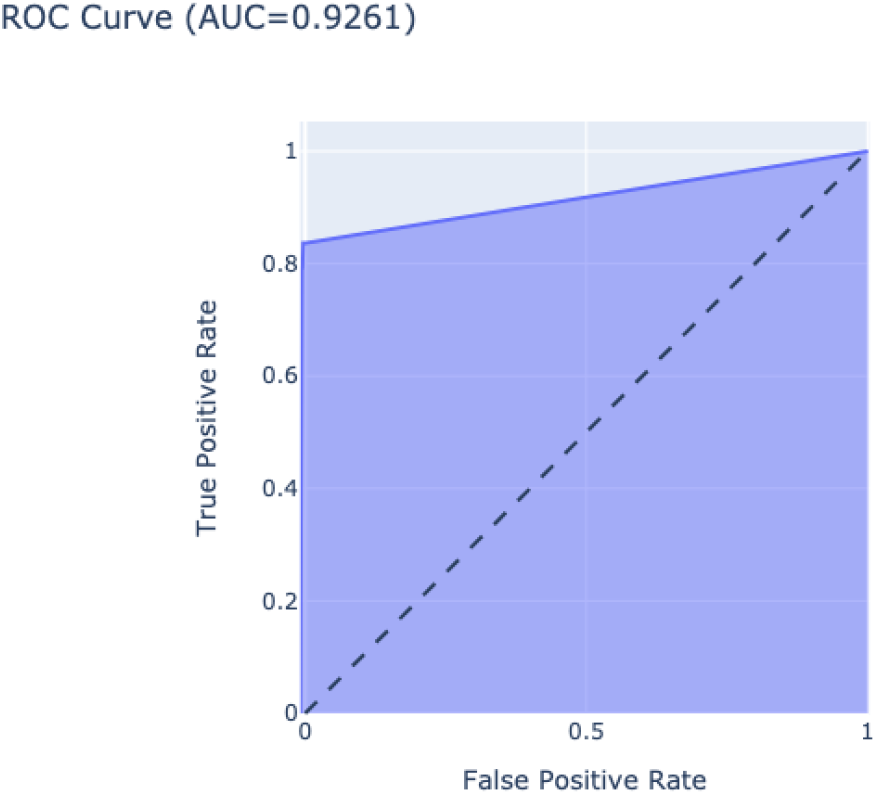
ROC curve for Rhodothermus profundi predictions.

### Test 2. Saccharomyces cerevisiae predictions

To predict the interactions of Saccharomyces cerevisiae, we considered Millerozyma farinosa and Naumovozyma dairenensis as similar species to train the model. Figure 9 displays the results obtained by Prots2Net. We can observe there that Prots2Net performed even better than in the previous test. Indeed, all the error metrics (accuracy, precision, sensitivity, specificity, MCC) reached values above 0.9, except in the very extreme values of the threshold. Such a good behavior is also reflected in Figure 10,where we show the error measures at a threshold of 0.5. We can see that only 6 interactions were wrongly predicted by Prots2Net. Also, we observe again the good performance of Prots2Net in Figure 11, where we display the *ROC* curve with a high *AUC* value of 0.9811.

**Figure 9.**
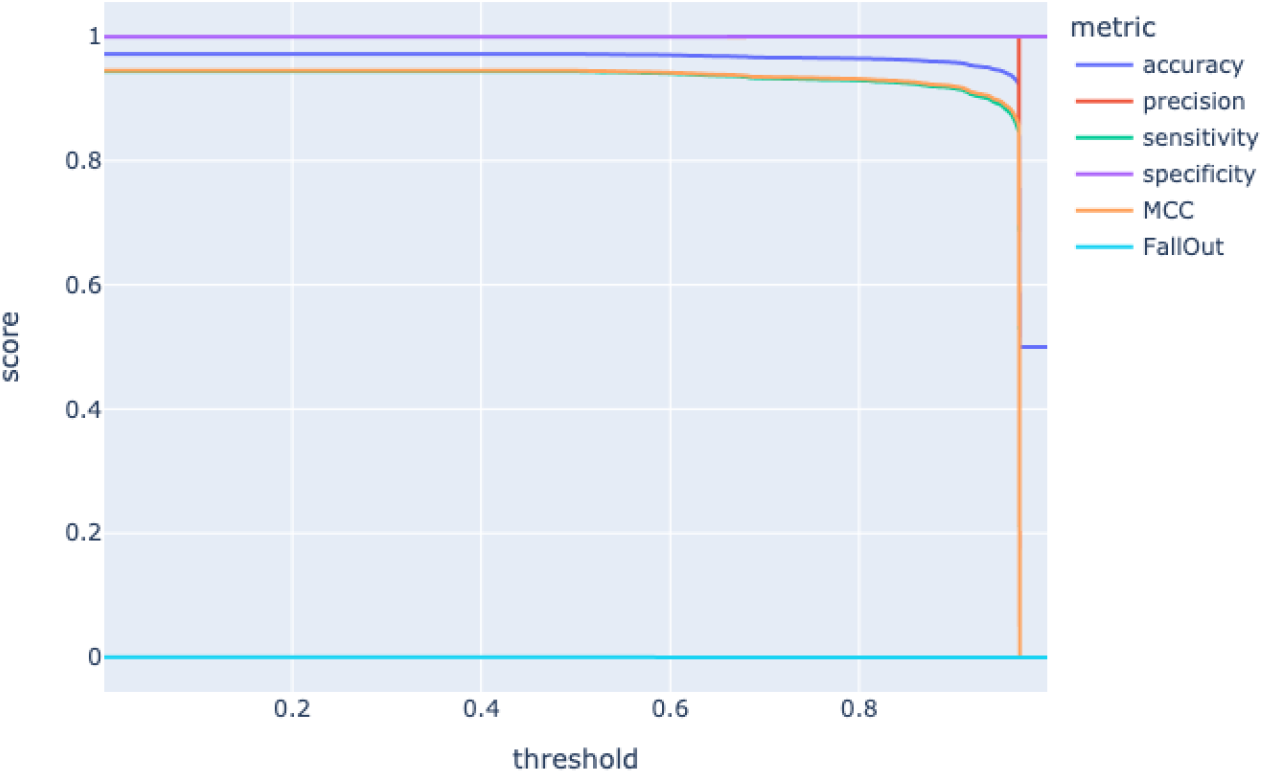
Metrics for Saccharomyces cerevisiae PPIs prediction.

**Figure 10.**
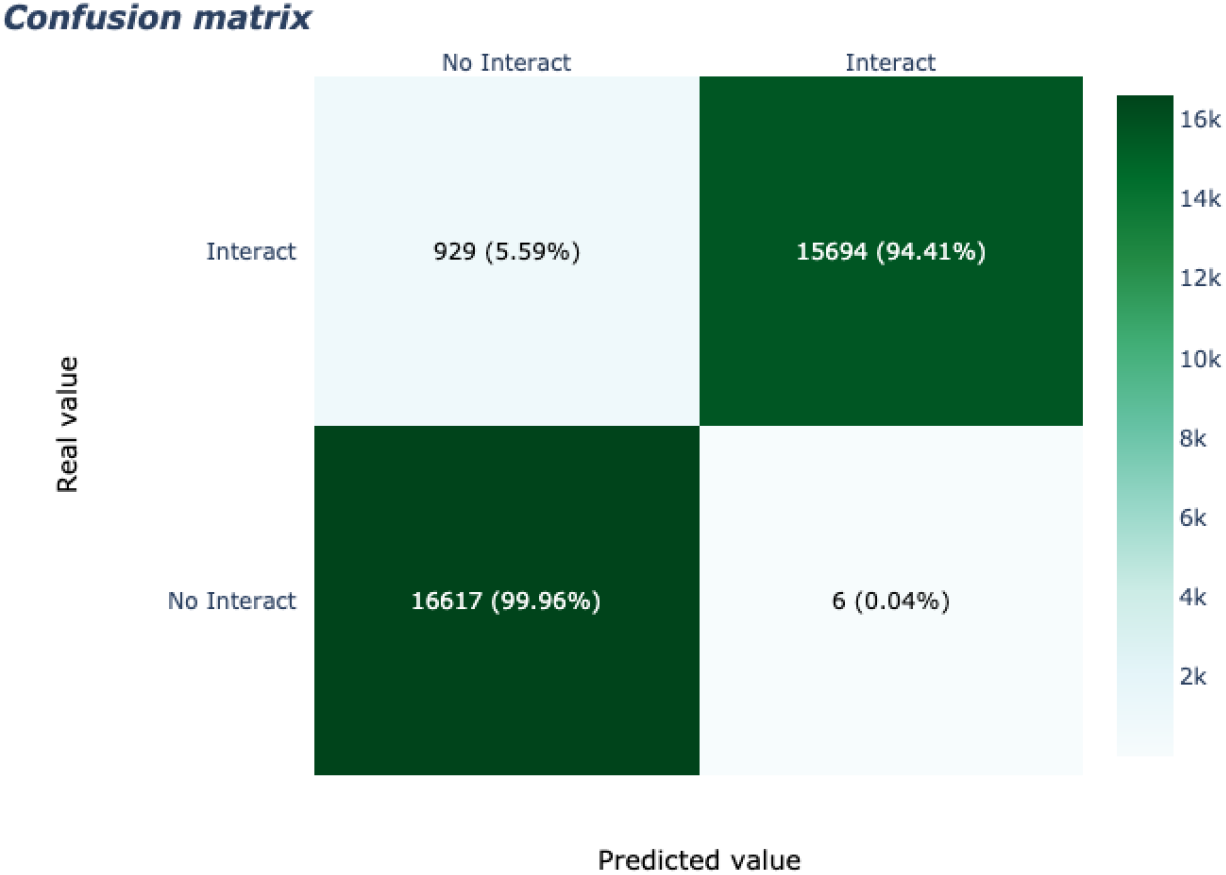
Confusion matrix for Saccharomyces cerevisiae with a threshold of 0.5.

**Figure 11.**
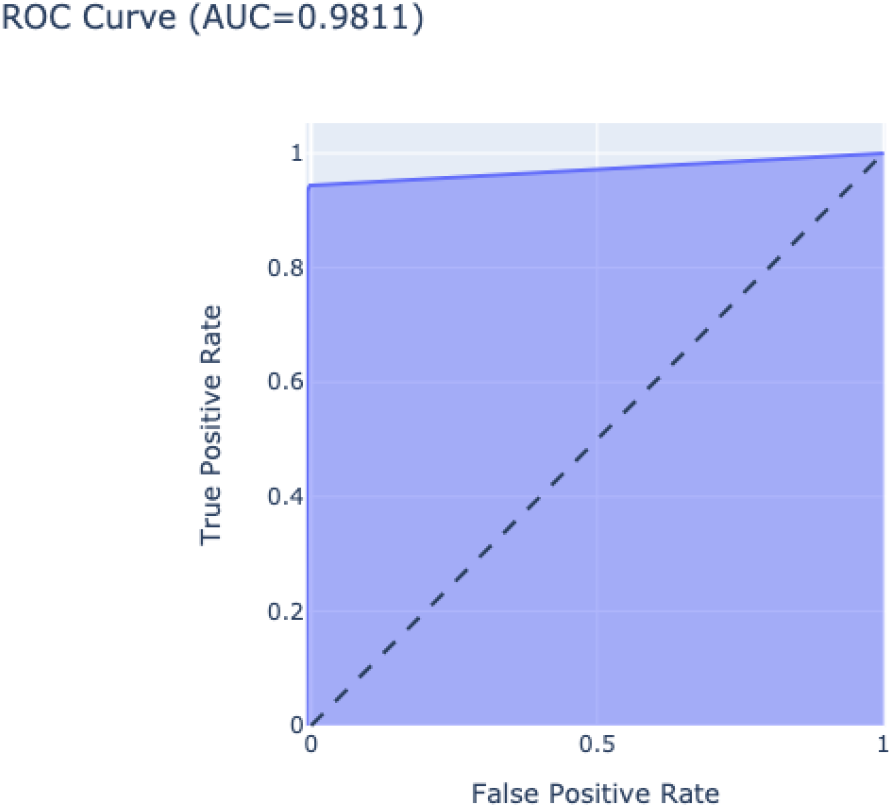
ROC curve for Saccharomyces cerevisiae PPIs prediction.

In order to contextualize the performance of Prots2Net in this test, in Table 1 we present the results of Prots2Net as well as the results of other 12 currently known methods (see the appendix section for a brief description of these methods). The evaluation measures of the other methods were obtained from [16]. We observe that Prots2Net has the best error measures. Hence, Prots2Net outperform all the other methods.

**Table 1.** Performance comparisons of 13 methods on the Yeast dataset

| Method | Acc | Sen | Pre | MCC |
| --- | --- | --- | --- | --- |
| Prots2Net | 97.18 | 94.41 | 99.96 | 94.52 |
| DeepPPI [4] | 94.43 | 92.06 | 96.65 | 88.97 |
| RF + LPQ [28] | 93.92 | 91.10 | 96.45 | 88.56 |
| Bio2Vec [27] | 93.30 | 92.70 | 93.55 | 87.49 |
| MCD + SVM [31] | 91.36 | 90.67 | 91.94 | 84.21 |
| LCPSSMMF [1] | 90.48 | 90.26 | 90.58 | 82.84 |
| 3-mers [27] | 90.26 | 88.14 | 91.65 | 82.38 |
| ACC [8] | 89.33 | 89.93 | 88.87 | N/A |
| LD [33] | 88.56 | 87.37 | 89.5 | 77.15 |
| AC [8] | 87.36 | 87.30 | 87.82 | N/A |
| PCA-EELM [32] | 87.00 | 86.15 | 87.59 | 77.36 |
| LD+KNN [30] | 86.15 | 81.03 | 90.24 | N/A |
| OLPP+RF [16] | 90.07 | 89.83 | 90.24 | 82.10 |

### Test 3. Homo sapiens predictions

To predict the interactions of Homo sapiens, we considered Pan paniscus and Gorilla gorilla as similar species to train the model. The performance of Prots2Net in this test is extraordinary, as we can observe in Figure 14. All the metrics are almost 1, except for the extreme values of the threshold. As remarkable, we can see in Figure 13, that with a threshold value of 0.5, Prots2Net miss 214 interactions, but all interactions are correctly predicted, i.e., there are no false positives. All these results are summarized in Figure 12, where the ROC curve has a *AUC* value of 0.9984.

**Figure 12.**
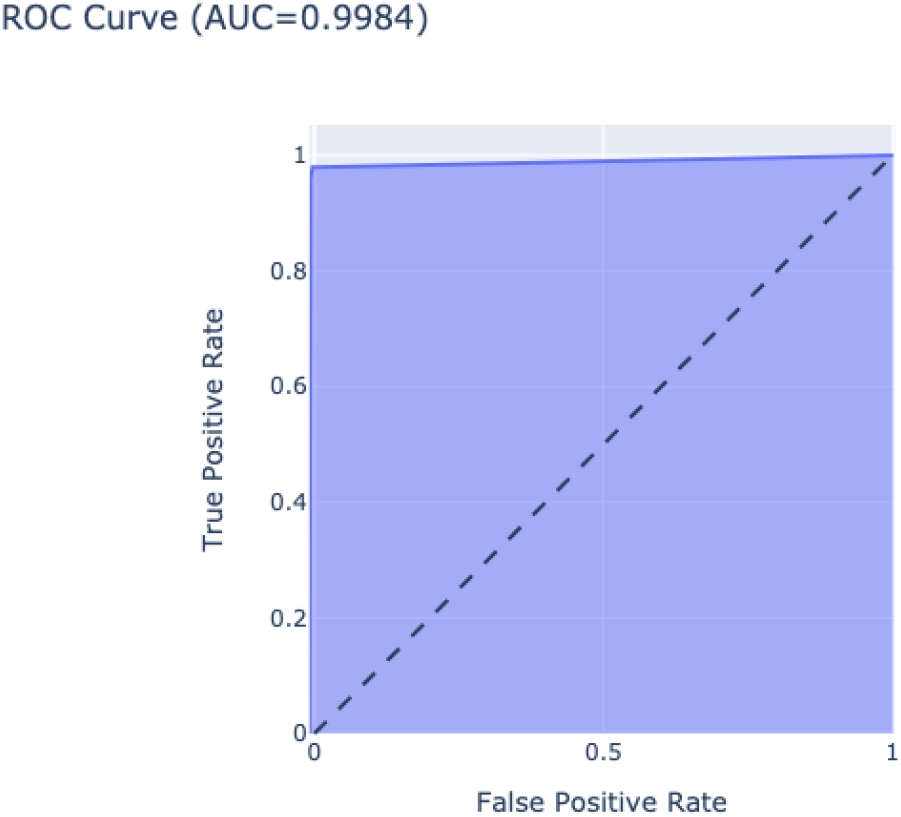
ROC curve for Homo sapiens PPIs prediction.

**Figure 13.**
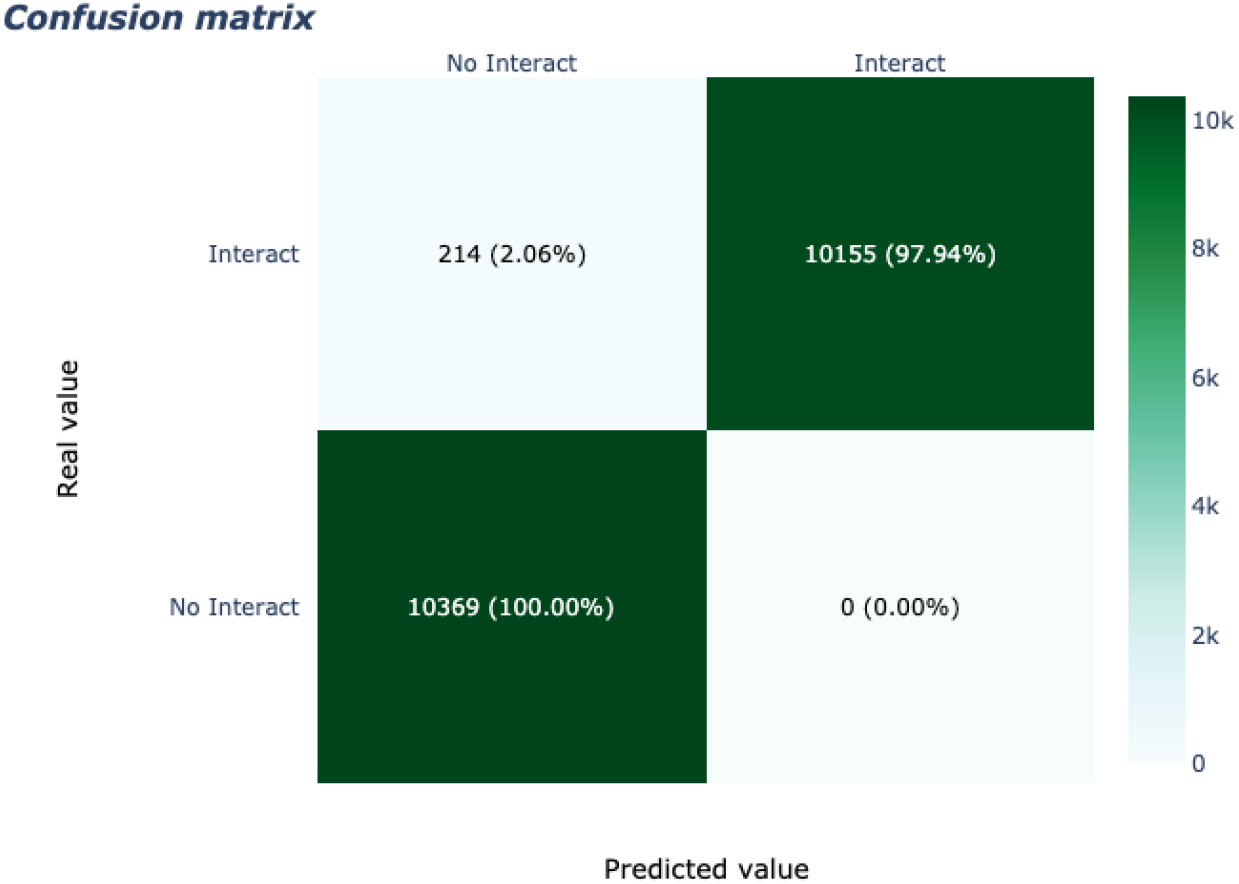
Confusion matrix for Homo sapiens with a threshold of 0.5.

**Figure 14.**
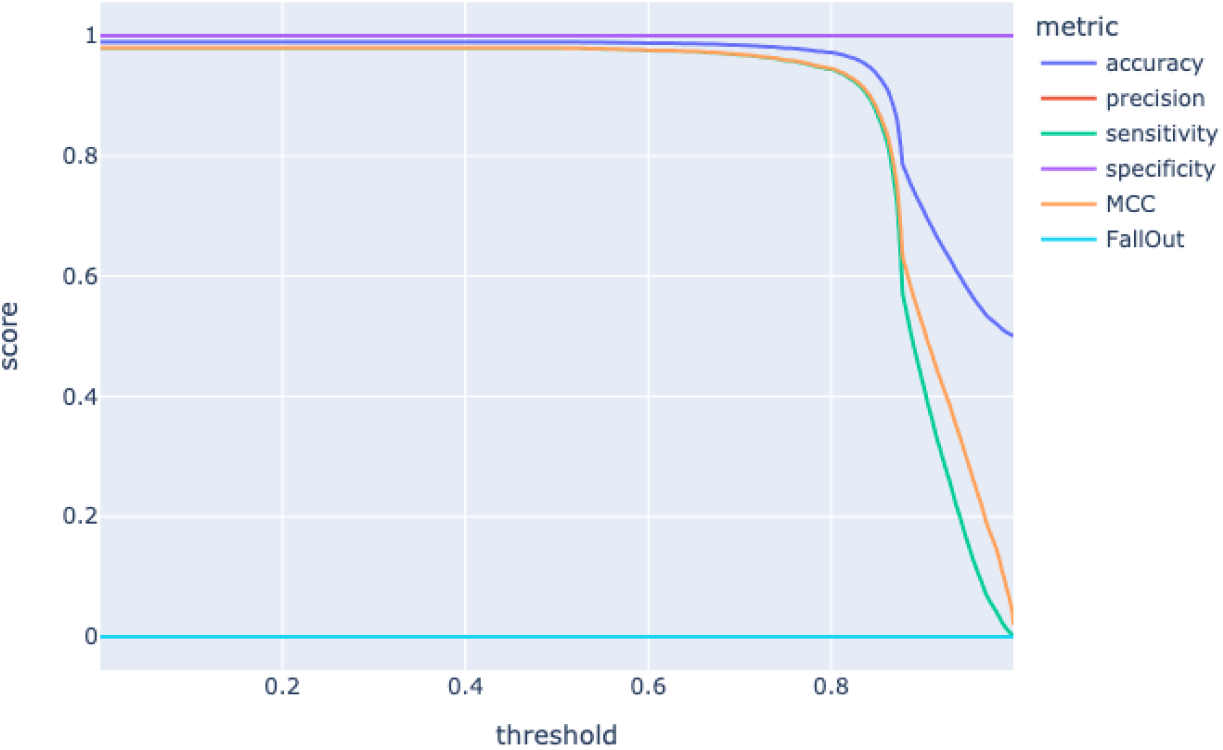
Metrics for Homo sapiens PPIs prediction.

As in the Yeast database, in order to compare the results obtained by Prots2Net with those obtained by other currently known methods, in Table 2 we present their evaluation measures. The evaluation measures of the other methods were obtained from [16]. We again can observe that Prots2Net obtained the best error measures. Hence, we conclude that Prots2Net out-perform all the other methods.

**Table 2.** Performance comparisons of 16 methods on the Human dataset.

| Method | Acc | Sen | Pre | MCC |
| --- | --- | --- | --- | --- |
| Prots2Net | 98.97 | 97.94 | 100 | 97.96 |
| DeepPPI [4] | 98.14 | 96.95 | 99.13 | 96.29 |
| MMI + NMBAC [3] | 97.57 | 96.57 | 98.30 | 95.13 |
| LDA + RF [20] | 96.4 | 94.2 | N/A | 92.8 |
| DCT + SMR + WSRC [11] | 96.30 | 92.63 | 99.59 | 92.82 |
| MMI [3] | 96.08 | 95.05 | 96.97 | 92.17 |
| LDA + RoF [20] | 95.7 | 97.6 | N/A | 91.8 |
| NMBAC [3] | 95.59 | 94.06 | 96.94 | 91.21 |
| AC + RF [20] | 95.5 | 94.0 | N/A | 91.4 |
| AC + RoF [20] | 95.1 | 93.3 | N/A | 91.0 |
| LDA + SVM [20] | 90.7 | 89.7 | N/A | 81.3 |
| AC + SVM [20] | 89.3 | 94.0 | N/A | 79.2 |
| PseAAC + RF [20] | 95.6 | 94.1 | N/A | 91.2 |
| PseAAC + RoF [20] | 95.3 | 93.6 | N/A | 90.7 |
| PseAAC + SVM [20] | 91.2 | 89.9 | N/A | 92.0 |
| OLPP+RF [16] | 96.09 | 95.20 | 96.56 | 92.47 |

### Impact of species selection

In order to test whether the selection of the two similar species affects the prediction results, we again considered predicting the PPIs in Homo sapiens but, with different pairs of similar species to train the model. Thus, we selected the following pairs of species.

1. Pan paniscus and Gorilla gorilla,
2. Arabidopsis thaliana and Arabidopsis lyrata.
3. Millerozyma farinosa and Naumovozyma dairenensis,
4. Rhodothermus profundi marinus and Salinibacter ruber,
5. Archaeglobus fulgidus and Methanopyrus kandlery AV19.

Notice that, we considered pairs of species at different taxonomy distance with Homo sapiens. As a first pair we consider two primates, as a second one two plants, followed by two fungi and then two bacteria and two archaea. As one would expect, the best results were obtained with the two primates, followed by the two plants and the two fungi. The worst results were obtained with the two bacteria and archaea. Figure 15 shows the error measures obtained in every test when considering different pairs of similar species.

**Figure 15.**
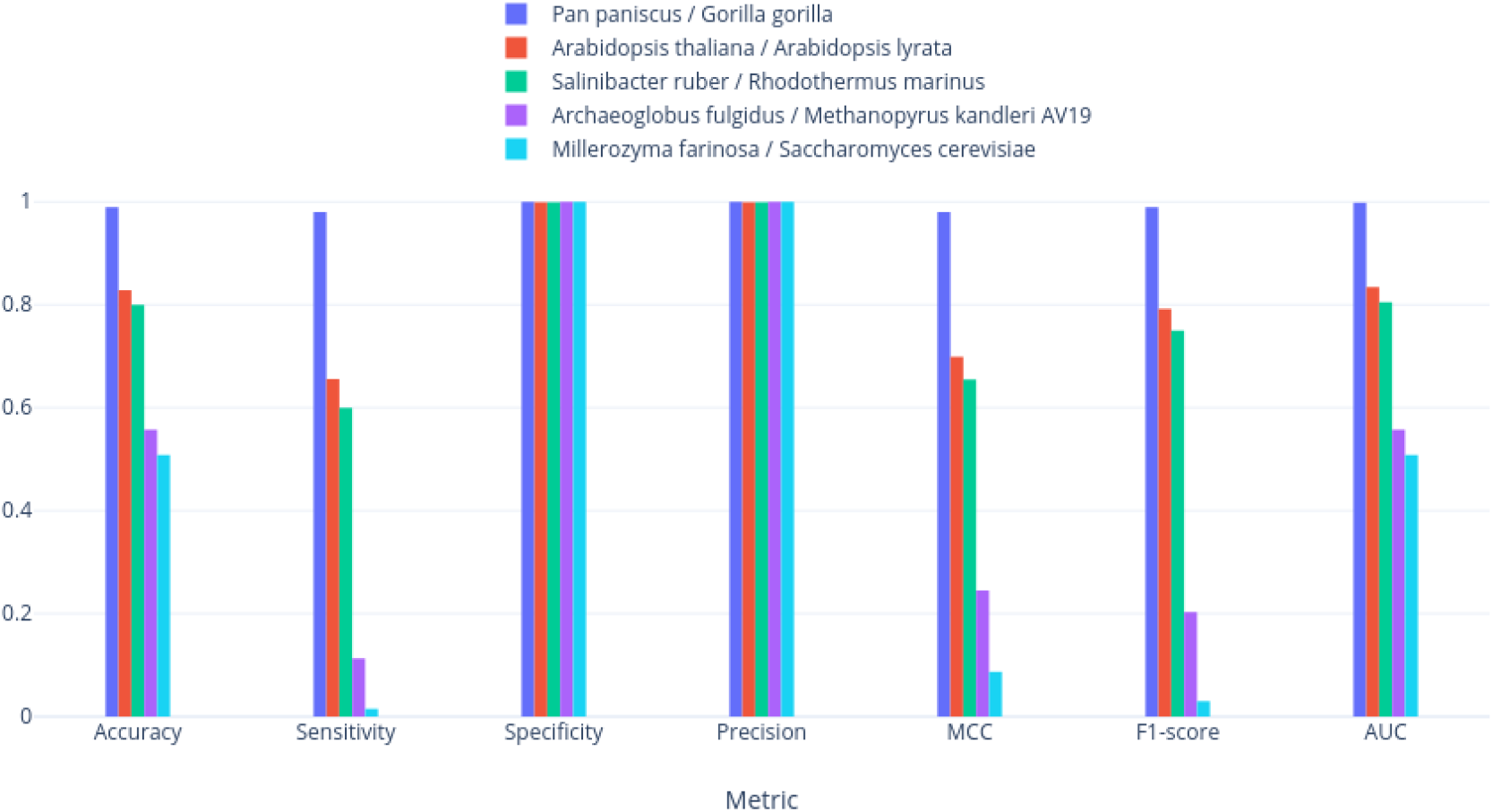
Metrics comparison to analyze the impact of the species selection.

We can observe there that, indeed, the best results were in the selection of two primates. We obtained a Specificity and Precision values of 1 in every pair of similar species. Recall that the Specificity is defined by 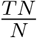 and the Precision is defined by 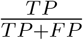. Also, we trained the model considering that two proteins interact when their combined interaction score in the STRING database is greater than 999. Hence, this high threshold ensures that the probability to truly interact when the model predicts an interaction is very high. Or, equivalently, there are very few false positives. On the other side, the other error measures clearly show that the model performed better with those pairs of PPINs that are taxonomically close to Homo sapiens.

## Conclusion

In this work, we present Prots2Net, a multilayer perceptron neural network designed to predict the PPIs of an input set of proteins. Our model takes into account sequence information only plus the PPI information available in the STRING database. The training data consists of two selected PPINs from the STRING database, which avoids the lack of a priori information needed to train the model.

The tests performed and reported in this paper to evaluate our tool, show that Prots2Net obtains very good results in the six error measures considered here. In addition, in the test presented here to evaluate the impact of the species selection, we conclude that the results improve when the selected species are taxonomically close to the species whose interactions are requested.

Regarding the comparison of Prots2Net with other existing methods, we conclude that Prots2Net outperform the other existing methods, in the yeast and human datasets, since it obtains the highest values of accuracy, sensitivity, precision, and Mathews correlation coefficient (MCC). Therefore, to consider the information of PPI data available on the STRING database clearly improves the PPI prediction. Hence, Prots2Net is an efficient tool to correctly predict PPIs from a metaproteome or a proteome sample with the advantage that, from the user’s point of view, only the file with the proteins sequences must be uploaded.

## Acknowledgements

We thank Jairo Rocha for useful discussions.

## Funding

This work has been supported by the Ministerio de Ciencia e Innovación (MCI), the Agencia Estatal de Investigación (AEI) and the European Regional Development Funds (ERDF) for its support to the project PGC2018-096956-B-C43.

## A PPIs prediction methods

We briefly describe here the PPIs prediction methods considered in the results and discussion section.

### A.1 DeepPPI

Deep neural networks for Protein–Protein Interactions prediction (DeepPPI) [4], employs deep neural networks to effectively learn representations of proteins from common protein descriptors. The framework of this algorithm consists on five steps:

1. Obtain protein interaction data from the DIP public database.
2. Obtain the protein sequences from PIR or UniProt
3. High quality features of protein sequence are extracted.
4. Obtain negative samples by random matching of protein in different subcellular locations.
5. Create training and test sets from the positive and negative samples, and feed a training set into the deep neural network.

### A.2 Rotation Forest & Local Phase Quantization

The method proposed in [28] use a Rotation Forest (RoF) classifier and the Local Phase Quantization (LPQ) descriptor from the Physicochemical Property Response (PR) Matrix of protein amino acids.

The RoF classifier [23] is a method for generating classifier ensembles based on feature extraction. To create the training data for a base classifier, the feature set is randomly split into K subsets (K is a parameter of the algorithm) and Principal Component Analysis (PCA) is applied to each subset in order to reduce its dimension.

The Local Phase Quantization (LPQ) [18] method is a common and efficient texture descriptor that adopts the Fourier transform to analyze the information in a matrix.

To borrow the feature extraction techniques from image processing, it is necessary to preprocess each amino acid sequence by transforming them into a matrix. The method, named Physicochemical Property Response Matrix (PR) [18], is used to represent the protein sequence.

### A.3 Bio2Vec

The bio2Vec [27] model was constructed based on a feature representation method for biological sequences called bio-to-vector (Bio2Vec) and a convolution neural network (CNN). The Bio2Vec obtains protein sequence features by using a “bio-word” segmentation system and a word representation model used for learning the distributed representation for each “bio-word”. Bio2Vec workflow consists of two stages:

1. Generate a fixed-length feature representation for each protein sequence. First segment the protein sequences into protein words, and then the protein words were transformed to a vector by Skip-Gram model.
2. Use a convolutional neural network with multiple convolution kernels for predicting PPIs. Given a pair of protein sequences, represent them using Bio2Vec and then concatenate them to form a feature pair.

### A.4 3-mers

In [27] a different bio word segmentation system is compared to Bio2Vec, consisting on split a sequence through a sliding window with stride s, where K is the size of window, in particular the 3-mers bio word segmentation is tested.

### A.5 MCD + SVM

The model presented in [31] is a combination of Multi-scale Continuous and Discontinous (MCD) feature representation and Support Vector Machine (SVM). On this model, the feature selection employed to construct an optimized and more discriminative feature set by excluding redundand features was mRMR. The workflow to predict the PPIs consist of three steps:

1. Represent protein sequences as a vector by using the proposed multi-scale continuous and discontinuous (MCD) feature representation
2. Minimum redundancy maximum relevance (mRMR) is utilized to do the feature selection
3. SVM predictor is used to perform the protein interaction prediction tasks.

### A.6 LCPSSMMF

LCPSSMMF [1] is a sequence-based feature extraction which combine local coding position-specific scoring matrix (PSSM) with multifeature fusion. The workflow of this model consist of three main steps:

1. Use a local coding method based on PSM to build PSSM (CPSSM), which incorporates global and local feature extraction to account for the interactions between residues in both continuous and discontinuous regions of amino acid sequences.
2. Adopt 2 different feature extraction methods (LAG and BP) to capture multiple key feature information by using the evolutionary information embedded in CPSSM.
3. Acquire feature vectors using the multifeature fusion method.

### A.7 AC and ACC

In [8] a method combining feature representation using auto cross covariance (ACC) and support vector machine (SVM) is present. AC accounts for the interactions between residues a certain distance apart in the sequence, so this method adequately takes the neighboring effect into account. This method consist of three main steps:

1. The amino acid residues are translated into numerical values representing physicochemical properties.
2. The numerical sequences are analyzed by ACC based on the calculation of covariance.
3. The SVM model is constructed using the vectors of AC variables as input.

ACC results in two kinds of variables, AC between the same descriptor, and cross covariance (CC) between two different descriptors. In [8] is analyzed the results to use only the AC variables (AC) and the AC plus CC variables (ACC).

### A.8 LD

In [33] a representation of local protein sequence descriptors and support vector machine (SVM) are combined. To represent the protein sequence, the following steps are done:

1. For each protein sequence, every amino acid is replaced by the index depending on its grouping.
2. Split the amino acid sequences into ten local regions of varying length and composition to describe multiple overlapping continuous and discontinuous interaction patterns within a protein sequence.
3. For each local region, three local descriptors, composition (C), transition (T) and distribution (D), are calculated.
4. The amino acids are divided into seven groups.
5. The descriptors for all local regions were combined, representing the general characteristics of the protein sequence.
6. The vector representing each protein pair is used as a feature vector for input into SVM.

### A.9 PCA-EELM

In [32] a method ensambling ELM and PCA is presented. The method workflow is:

1. Four kinds of useful sequence-based features such as Auto Covariance (AC), Conjoint triad (CT), Local descriptor (LD) and Moran autocorrelation (MAC) are extracted from each protein sequence to mine the interaction information in the sequence.
2. A feature reduction method PCA is employed to extract the most discriminative new feature subset.
3. ELM classifier is constructed using the vectors of resulting feature subset as input.

### A.10 LD + KNN

In [30] a method using local descriptors and k-nearest neighbors (KNNs) learning system is presented with the following workflow:

1. Represent each protein sequence as a vector using a local protein sequence descriptors
2. Characterize a protein pair in different feature vectors by coding the vectors of two proteins in this protein pair
3. Construct a KNNs model using the feature vectors of the protein pair as input

### A.11 OLPP + RF

In [16] a computational method for predicting PPIs based on combining orthogonal locality preserving projections (OLPP) and rotation forest (RoF) models is presented. The method consist of the following steps:

1. Convert the protein sequence into position-specific scoring matrices (PSSMs) containing protein evolutionary information by using the Position-Specific Iterated Basic Local Alignment Search Tool (PSI-BLAST)
2. Characterize the proteins as a fixed length feature vector by applying OLPP to PSSMs
3. Train the RoF classifier.

### A.12 MMI + NMBAC

In [3] two different methods to extract features from protein sequence information are used:

- Use k-gram feature representation calculated as Multivariate Mutual Information (MMI)
- Extract additional feature by normalized Moreau-Broto Autocorrelation (NMBAC)

These two approaches are used independently or combined to transfrom protein sequence into feature vectors. Then a Random Forest model is feed with this feature vectors to distinguish interaction pairs from non-interaction pairs.

### A.13 PseAAC — AC — LDA + RF — RoF — SVM

In [20] three different feature extraction methods are used:

- Pseudo amino acid composition (PseeAAC) that calculates the correlations between residues in the whole protein sequence.
- Auto Covariance (AC) that takes the neighboring effect into account and describes the average interactions between residues with a certain distance apart.
- Latent dirichlet allocation (LDA) which is a generative probabilistic model.

Then these methods are combined with these three classifiers

- Support Vector Machine (SVM)
- Random Forest (RF)
- Rotation Forest (RoF)

In [20] the proposed model is the method that combines the generative LDA and discriminative RF models.

### A.14 DCT + SMR + WSRC

The method proposed in [11] consists on using discrete cosine transform (DCT) on substitution matrix representation (SMR) and then use a weighted sparse representation based classifier (WSRC), a variant of traditional SRC, which integrates both sparsity and locality structure data.

Discrete cosine transform is a popular linear separable transformation in the lossy signal and image compression processing for its powerful energy compaction property.

In SMR, a *N ×* 20 matrix is generated to represent a *N* ′ length protein sequence based on a substitution matrix.

## Notes

https://github.com/adriaalcala/prots2net

### Competing Interest Statement

The authors have declared no competing interest.

